# Transcriptional profiling of *Drosophila* male-specific P1 (pC1) neurons

**DOI:** 10.1101/2023.11.07.566045

**Authors:** Osama M. Ahmed, Amanda Crocker, Mala Murthy

## Abstract

In *Drosophila melanogaster*, the P1 (pC1) cluster of male-specific neurons both integrates sensory cues and drives or modulates behavioral programs such as courtship, in addition to contributing to a social arousal state. The behavioral function of these neurons is linked to the genes they express, which underpin their capacity for synaptic signaling, neuromodulation, and physiology. Yet, P1 (pC1) neurons have not been fully characterized at the transcriptome level. Moreover, it is unknown how the molecular landscape of P1 (pC1) neurons acutely changes after flies engage in social behaviors, where baseline P1 (pC1) neural activity is expected to increase. To address these two gaps, we use single cell-type RNA sequencing to profile and compare the transcriptomes of P1 (pC1) neurons harvested from socially paired versus solitary male flies. Compared to control transcriptome datasets, we find that P1 (pC1) neurons are enriched in 2,665 genes, including those encoding receptors, neuropeptides, and cell-adhesion molecules (*dprs*/*DIPs*). Furthermore, courtship is characterized by changes in *∼*300 genes, including those previously implicated in regulating behavior (e.g. *DopEcR, Oct*β*3R, Fife, kairos, rad*). Finally, we identify a suite of genes that link conspecific courtship with the innate immune system. Together, these data serve as a molecular map for future studies of an important set of higher-order and sexually-dimorphic neurons.

## Introduction

*Drosophila melanogaster* male flies court conspecific female flies by displaying time-varying behaviors comprising tapping with the foreleg tarsi, chasing, singing via unilateral wing vibrations, and abdominal bending (1, 2). These complex, male-typical behaviors depend on a developmental program that is largely orchestrated by sex-specific expression of the transcription factor genes *fruitless* and *doublesex* (3–9). Sex-specific expression of *fru* and *dsx* is required for the proper development of a cluster of 20 interneurons per hemibrain, called “P1 (pC1)” (also known as pMP4/e (10, 11)), that is critical for the display of male-typical courtship behavior (3, 7, 12–14). P1 (pC1) neurons have extensive dendritic arborization in the lateral protocerebrum (10, 11), where they integrate visual, olfactory, and gustatory information to control courtship decisions (14–17), and their neuronal activity is modulated by social experience (13, 14, 18, 19). Inhibition of P1 (pC1) neurons precludes wildtype male courtship, while their activation elicits courtship behavior, even in the absence of a mating partner (3, 12, 14, 17). These neurons also regulate courtship behaviors at multiple timescales. At short timescales, P1 (pC1) neurons disinhibit courtship song-circuitry to complexify song bouts on a moment-to-moment basis (20), and P1 (pC1) activity continuously sensitizes LC10a visual projection neurons to tune the vigor of courtship chasing (21). At longer timescales, P1 (pC1) neurons set a courtship arousal state (22, 23) that is modulated by dopamine (18) and decreases with successive matings (24). The role of P1 (pC1) neurons, however, is not limited to driving courtship behaviors (25).

At low activation levels, P1 (pC1) neurons promote male-male aggression (23, 26). Moreover, activation of P1 (pC1) neurons, either experimentally (13, 23) or via conspecific contact (14–16, 27), generates a minutes-long social arousal state that sensitizes male flies to both courtship and aggression, depending on social context (28). Beyond mating and fighting, P1 (pC1) neurons also balance other competing drives, including sleeping and feeding; P1 (pC1) neurons contribute to sleep homeostasis (29) and sleep-need (30, 31), and the presence of high quality foods or starvation leads to a net inhibition of P1 (pC1) neurons, such that hungry flies prioritize feeding over mating (32, 33). Repeated matings also diminish dopamine signals that gate P1 (pC1)’s ability to drive courtship, effectively shunting everlasting courtship attempts (18). Balancing which drive (feeding, sleeping, mating, etc.) takes precedence in any given moment depends not only on dopamine-, but on insulin, octopamine-, and tyramine-mediated signaling onto P1 (pC1) neurons. Therefore, for example, knowing the receptor expression of P1 (pC1) neurons will clarify how these important neurons are modulated.

Despite the role of P1 (pC1) neurons in regulating motivated behaviors, we know little about the molecular profile of these neurons, or how that profile changes with behavior. Addressing these gaps is critical for understanding how neuromodulation, synapse function, and signaling pathways contribute to P1 (pC1)’s diverse functions, including their role in courtship, across different timescales. That is, we may better understand how P1 (pC1) neurons influence internal states and distinct behaviors by resolving their molecular profiles. For example, different neuromodulatory signals may alter P1 (pC1) physiology in distinct ways; therefore, knowing which receptors are expressed in P1 (pC1) neurons will clarify which signals they are sensitive to, and on which timescales. P1 (pC1) neurons themselves may push and pull on downstream neurons via distinct neuromodulatory pathways (34), yet we do not know the full suite of neuromodulators and neuropeptides that these neurons produce. Importantly, P1 (pC1) neuronal activity is shaped directly by social interactions (14–16, 18, 21), suggesting that social interactions dynamically alter P1 (pC1)’s transcriptional milieu. To address these gaps, we resolved P1 (pC1)’s transcriptome via single cell-type RNA sequencing and found 2,665 differentially-expressed genes in P1 (pC1) neurons compared to control whole-head and whole-brain datasets. We also found that the gene expression profile of P1 (pC1) neurons harvested from male flies paired with conspecific female flies, where we expect increased P1 (pC1) neural activity due to direct social contact (14, 18, 21), is distinct from solitary, unpaired male flies, with 322 genes showing differential expression. By comparing these 322 genes with extant datasets, we identified a common set of genes, including many immune-response genes, that are linked to conspecific courtship. In total, our results provide a basis for understanding how the transcriptional profile of P1 (pC1) neurons contributes to their important function as higher-order regulators of complex and distinct behaviors.

## Results

### Single cell-type RNA sequencing of P1 (pC1) neurons

There are ∼2,000 fruitless+ neurons in the central brain (6, 8, 11, 37), ∼40 of which form the P1 (pC1) clusters (∼0.02% of all central brain neurons) (Figure 1A). We labeled P1 (pC1) neurons with a fluorescent reporter (NP2631, fruFLP»GFP) (11), allowing us to reliably harvest 10-12 neurons per fly (Figure 1B); the average CPM (counts per million, log2 scale) for GFP was 5.235. We then assayed gene expression by sequencing RNA from neurons harvested from male flies that were either socially isolated (n = 4 unpaired) or paired (n = 4) with a mated female conspecific. We pooled neurons for each male fly and amplified mRNA by using the poly-T primer-based SMARTer kit. We then sequenced cDNA, at a depth of 30 million reads. We found 2,665 genes that were upregulated in P1 (pC1) neurons relative to our control datasets (Figure 1C), with 994 of these being currently unidentified genes (CG genes) and 1,509 genes with expected high expression in fly brains according to FlyAtlas AffyCall (via FlyMine) (38). Our control datasets comprised previously published whole-head male fly transcriptomes (35), in which “naive” male flies were either socially isolated (n = 4) or housed with a mated female conspecific (n = 4). For clarity, we refer to “naive” and “trained” datasets as “unpaired” and “paired” control datasets, respectively. Additionally, in order to test for sexually dimorphic gene expression and assess the quality of our transcriptomes, we generated whole-brain female fly transcriptomes (n = 2) alongside our P1 (pC1) neuron datasets (36).

**Figure 1.**
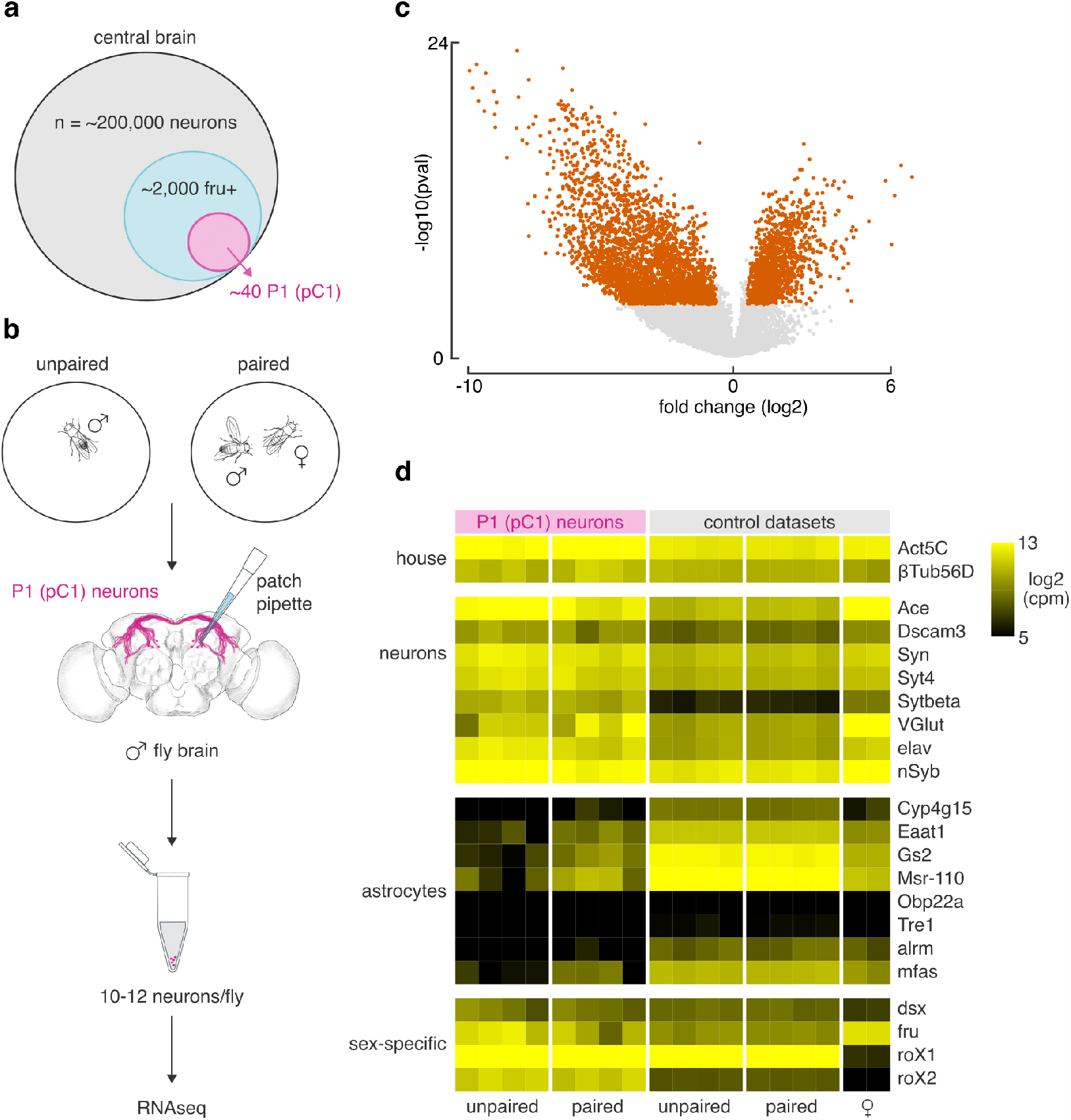
Single cell-type RNA sequencing of fru+ P1 (pC1) neurons. **(A)** P1 (pC1) neurons comprise roughly 2% of fruitless+ (fru+) neurons and 0.02% of all neurons in the central brain. **(B)** Schematic of courtship assay and protocol for harvesting P1 (pC1) neurons and RNA amplification. Male flies were isolated (unpaired) or placed with a conspecific female for 30 minutes (paired) prior to harvesting P1 (pC1) neurons (10-12 neurons per male). **(C)** Volcano plot of genes that are differentially expressed in P1 (pC1) neurons (orange). **(D)** Heatmap of gene expression. Each row denotes a gene with an expected expression pattern in P1 (pC1) neurons, organized by function: house-keeping genes, neuron-specific genes, astrocyte-specific genes, and sex-specific genes. Each column denotes an independent dataset. Unpaired and paired whole-head control datasets were previously published (35). The two female whole-brain control datasets were generated alongside P1 (pC1) neurons and previously published (36). cpm, counts per million.

As a quality check, we first assayed enrichment of gene transcripts that we expected to find in P1 (pC1) neurons. We found that housekeeping genes (*Act5c, βTub56D*) and neurontypical genes (*Ace, Dscam3, Syn, Syt4, Sytβ, VGlut, elav, nSyb*) were highly enriched in P1 (pC1) neurons, and, as expected, we found that astrocyte-typical genes (*Cyp4g15, Eaat1, Gs2, Msr-110, Obp22a, Tre1, alrm, mfas*) were expressed at low levels (Figure 1D). Additionally, we found that many stressrelated genes encoding heat shock proteins, with one exception (*Hsp70Bb*), were also expressed at low levels, suggesting that our experimental workflow was not unduly stressful to the animals (Figure S1). We also found enrichment of long noncoding RNAs (*roX1, roX2*) (39) and transcription factors (*dsx, fru*) that are expected to be found in male-specific neurons (Figure 1D). Collectively, these results boost our confidence that the P1 (pC1) transcriptome we generated is of high quality and not significantly contaminated by glial cells.

### Receptor gene expression of P1 (pC1) neurons

It is not fully known how P1 (pC1) neural activity is modulated by upstream partners. Dopamine, for example, acts through DopR2, which is expressed in P1 (pC1) neurons (Figure 2), to control mating drive (18). What other signals are P1 (pC1) neurons sensitive to? To answer this question, we determined which receptor-encoding genes are highly expressed in P1 (pC1) neurons. We found that P1 (pC1) neurons are enriched in 13 genes encoding neuromodulatory receptors that bind serotonin (5-HT), dopamine, octopamine, and tyramine (Figure 2). We were surprised to find enrichment of 6 sensory receptor genes, including 5 putative chemoreceptors (Or1, Or45a, Or63a, Ir56a, Gr28b). We previously uncovered these genes, with the exception of *Or1*, in at least one cell type of mushroom body neurons (36), suggesting that these genes are more broadly expressed than their names suggest. We found enrichment of several genes encoding for neuropeptide receptors (n = 11) and hormone receptors (n = 6), including the sex peptide receptor-encoding gene (*SPR*) and the activity-dependent immediate early gene *Hr38*, respectively. We also found enrichment of genes encoding canonical receptors for acetylcholine, glutamate, and GABA (Figure S2), including *Rdl* (adjusted p-value = 5.07e-21) which is known to be expressed in P1 (pC1) neurons (16).

**Figure 2.**
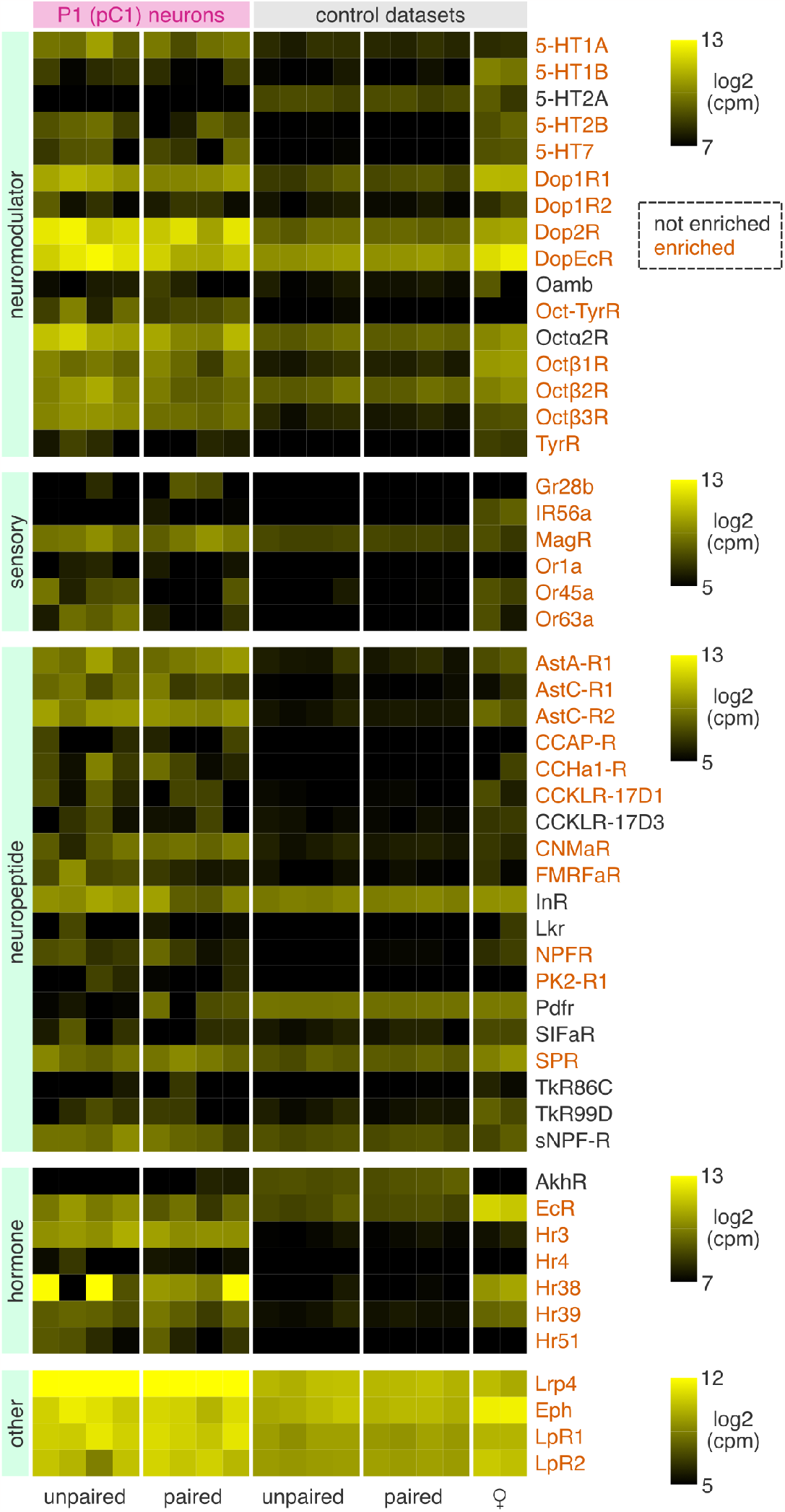
Receptor gene expression profile of P1 (pC1) neurons. Each row denotes a single receptor-encoding gene, organized by receptor type: neuromodulatory, sensory, neuropeptide, and hormone receptors, with 4 genes (*Lrp4, Eph, LpR1, LpR2*) categorized as “other”. Each column denotes an independent dataset (as in heatmap in Figure 1). Genes enriched in P1 (pC1) are annotated in orange.

We profiled 4 other receptors (*Lrp4, Eph, LpR1*, and *LpR2*), all of which were highly expressed in P1 (pC1) neurons. *Lrp4* (CG8909), which codes for a transmembrane protein, is the most enriched gene in P1 (pC1) neurons. *Lrp4* is expressed throughout the central brain of *Drosophila* where it controls synaptogenesis (40, 41) and regulates synapse number in excitatory neurons (40). While loss of *Lrp4* diminishes olfactory-driven attraction to apple cider vinegar, it is unknown if loss of *Lrp4* also impacts olfactory-driven social behaviors. Our results suggest that LRP4 may contribute to P1 (pC1) physiology and therefore a potential target for future study (41).

Neuropeptidergic and neuromodulatory signals dial P1 (pC1) neuron activity up or down to regulate behaviors (34, 42). For example, P1 (pC1) neurons do not appear to be directly postsynaptic to Drosulfakinin+ (DSK+) neurons, but could nonetheless be privy to DSK-mediated neuromodulation (43). DSK regulates male courtship behavior via its receptor CCKLR-17D3 (43) and aggression through its other receptor CCKLR-17D1 (44). We did not find that *CCKLR-17D3* is specifically enriched in P1 (pC1) neurons; however, *CCKLR17D1* is. This suggests that DSK may modulate P1 (pC1) neurons via distinct pathways. Additionally, neuromodulatory loops keep mating drive in check (18, 24, 45). For example, P1 (pC1) neurons excite male-specific NPF neurons, which then inhibit neurons that express the NPF receptor, NPFR, to diminish courtship drive (45). A parsimonious feedback model poses that P1 (pC1) neurons are themselves the directs targets of NPFmediated inhibition. However, P1 (pC1) neurons do not overlap with NPFR+ neurons (45), suggesting that P1 (pC1) neurons do not express *NPFR*. In contrast, our data show that *NPFR* is enriched in P1 (pC1) neurons. This difference may be due to technical distinctions in our approaches, but it suggests that P1 (pC1) neurons may be directly modulated by NPF. All together, mating and fighting may be differentially regulated at the level of P1 (pC1) via distinct neuropeptidergic and neuromodulatory signals, although we cannot discern if these differences arise from subsets of molecularly distinct neurons in the cluster.

P1 (pC1) neurons are inhibited by high-quality food and by starvation, which promotes feeding in lieu of courtship (32, 33). This inhibition is mediated by insulin (33) and tyramine (32) potentially acting on their cognate receptors, INR and TAR3 respectively. However, we did not find enrichment of *InR* (CG18402) or *Tar3* (CG16766) in P1 (pC1) neurons. *InR* is broadly-expressed throughout the nervous system (46) so we did not expect it to be specifically upregulated in P1 (pC1) neurons. As for *Tar3*, RNAi-mediated knockdown of TAR3 does not abolish P1 (pC1) neural responses to tyramine, nor does it abolish courtship in fed male flies (32). This suggests that other tyramine receptors may be expressed in P1 (pC1) neurons to regulate tyramine-mediated responses. Indeed, we do find enrichment of two other tyramine receptor-encoding genes *TyrR* (CG7431) and *Oct-TyrR* (CG7485) (Figure 2), suggesting these are potential targets for tyramine’s effect on P1 (pC1) neurons. Overall, P1 (pC1) neurons express many receptor genes, of various classes, including genes with known functional effects on P1 (pC1) physiology (e.g. *Rdl, DopR2*). Although differentially-expressed genes are typically translated into protein (47), it is possible that not all receptor genes translate into functional receptors. Regardless, our results suggest that P1 (pC1) neurons are broadly sensitive to myriad excitatory and inhibitory signals that act in concert to fine-tune P1 (pC1) activity.

### Neuropeptide gene expression of P1 (pC1) neurons

Neuropeptides regulate behavior (42). Which neuropeptides might P1 (pC1) neurons use to modulate the activity of downstream partners? To address this, we profiled the expression of 49 neuropeptide-encoding genes and found that 10 of these genes (*AstC, AstCC, DH31, Dh44, Gpb5, Ilp7, NPF, SP, sNPF, spab*) were highly enriched in P1 (pC1) neurons (Figure 3). Several of these genes are expressed in fruitless+ neurons or contribute to courtship behaviors. For example, *Dh31* and *Ilp7* are highly expressed in fruitless+ neurons of both sexes (48). And, Neuropeptide F (*NPF*) regulates courtship drive in male flies (45). P1 (pC1) neurons are expected to be cholinergic (30); we confirm this by finding high expression of *Ace, ChAT, VAChT*, and *ChT* in P1 (pC1) neurons (adjusted p-values = 1.7e-17, 9.98e-21, 2.54e-13, and 4.45e-39 respectively, Figure S3). Cholinergic neurons in the fly midbrain tend to express the neuropeptide-encoding genes *Tk, amn, sNPF*, and *spab* and not *PDF, Mip*, and *SIFa* (49). Our findings align with this; *sNPF* and *spab* are enriched while *PDF, Mip*, and *SIFa* are absent or expressed at low levels in P1 (pC1) neurons (Figure 3). *sNPF* and *spab* are expressed in 20-25% of midbrain neurons and are hypothesized to be modulatory co-transmitters (49). Additionally, *spab* is generally not co-expressed with *Nplp1* (49); our results also align with this finding—*spab*, but not *Nplp1*, is enriched in P1 (pC1) neurons. All together, these results give us confidence that we are capturing key neuropeptides expressed in P1 (pC1) neurons.

**Figure 3.**
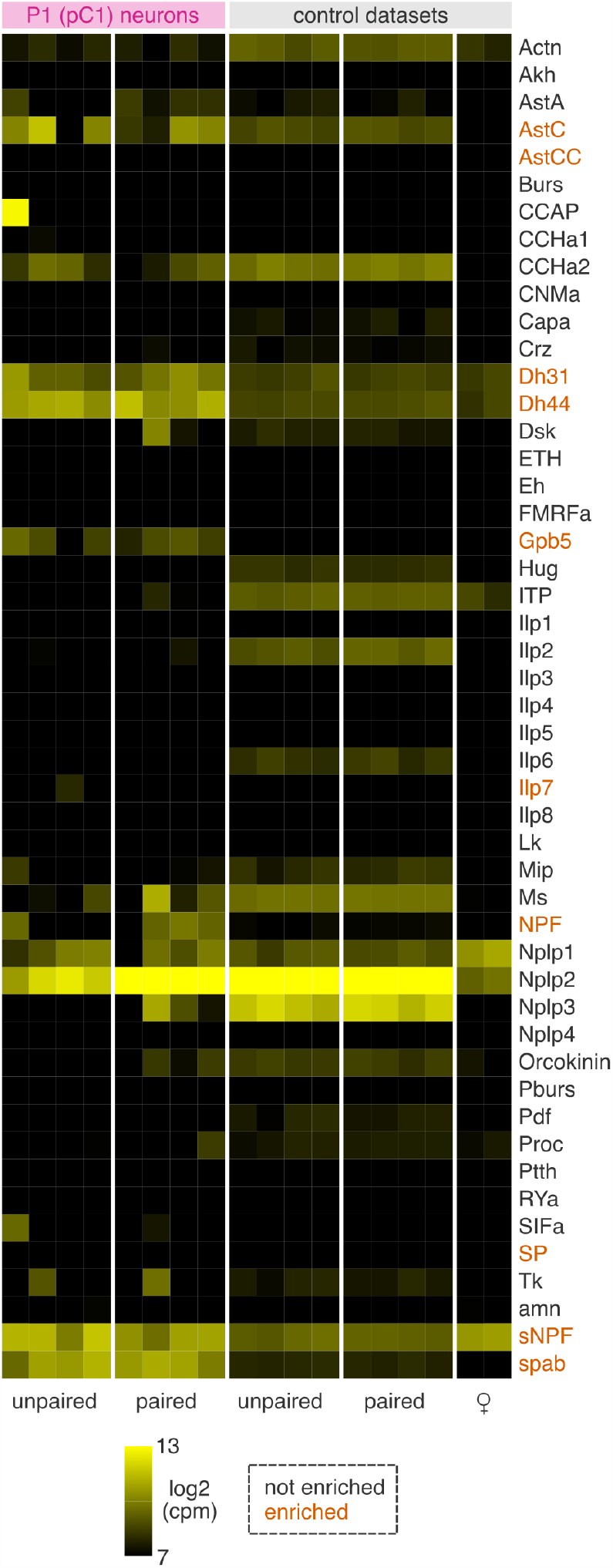
Neuropeptide gene expression profile of P1 (pC1) neurons. Each row denotes a single neuropeptide-encoding gene. Each column denotes an independent dataset (as in heatmap in Figure 1). Genes enriched in P1 (pC1) are annotated in orange.

We find that P1 (pC1) neurons express two genes, *AstC* and *AstCC*, that encode for allatostatin C-like peptides. *AstC* is expressed in a subset of clock neurons (50) including DN1 neurons; DN1 neurons coordinate with P1 (pC1) neurons to prioritize sleep or courtship depending on the animal’s needs (30). That *AstC* and *AstCC* have a functional role in P1 (pC1) neurons is yet to be determined. We were surprised to find enrichment of *Sex Peptide* (*SP*) in P1 (pC1) neurons (Figure 3). SP is produced by the male accessory glands and transferred to female flies during copulation (51). It acts through its high-affinity receptor SPR, which is broadly expressed in the nervous system of female flies (52), to regulate female post-mating behaviors (53). A role for SP in regulating central courtship circuitry of male flies is unknown. Finding *SP* enrichment in P1 (pC1) neurons suggests the exciting possibility that SP might have a functional role in regulating neural circuitry in both sexes.

The diuretic hormone Dh44 contributes to nutrient sensing (54), aggression (55), and stress responses (56). We find that P1 (pC1) neurons differentially express *Dh44*, in keeping with a recent analysis of fruitless+ neurons showing that *Dh44* is expressed in a population of fruitless+ cholinergic neurons (48). Moreover, the morphology of some of these neurons resembles P1 (pC1). We also find that P1 (pC1) express *Dh31*, a calcitonin ortholog (56) that contributes to locomotor activity in flies (57) and is widely expressed in the fly nervous system (58). There are ∼ 193 Dh31 neurons in the fly brain (58) and co-expression experiments will clarify if some of these neurons are indeed P1 (pC1). We also find enrichment of *Gpb5*, which encodes a glycoprotein hormone that contributes to reproduction in diverse species (59). Our results indicate that Dh44, Dh31, and Gpb5, which regulate distinct behaviors, are candidates for future studies of P1 (pC1) function. Overall, we have generated a short list of neuropeptides that may contribute to P1 (pC1)’s diverse functional roles in regulating behavior.

### Comparison of P1 (pC1) and fruitless+ neuronal gene expression

P1 (pC1) neurons are a subset of fruitless+ neurons and they therefore likely express a suite of genes that is common to this population. To test this, we cross-checked the list of 2,665 genes that are upregulated in P1 (pC1) neurons with a recently-generated transcriptome of fruitless-expressing neurons (60). We find that almost half (n = 1,326) of the genes in our list are present in this dataset (Figure 4A). Most of these genes are typically upregulated in brain tissue (Figure 4B) and downregulated in the gut and testis (Figure 4C) suggesting that these genes primarily underpin neuron-typical functions. We test this idea by performing Gene Ontology (GO) enrichment analysis on the 1,326 genes. We find that many genes map to GO terms related to biological regulation (n = 668 genes, p-value = 3.28e-33), cell communication (n = 341 genes, p-value = 2.41e-27), and synapse organization (n = 108 genes, p-value = 6.85e-28) among others, suggesting that most of the overlapping genes contribute to neuronal function. We also find that the top hit for protein domain enrichment comprises genes in the immunoglobulin-like domain superfamily (n = 57 “IgSF” genes of the 1,326 genes, p-value = 4.63e-113).

**Figure 4.**
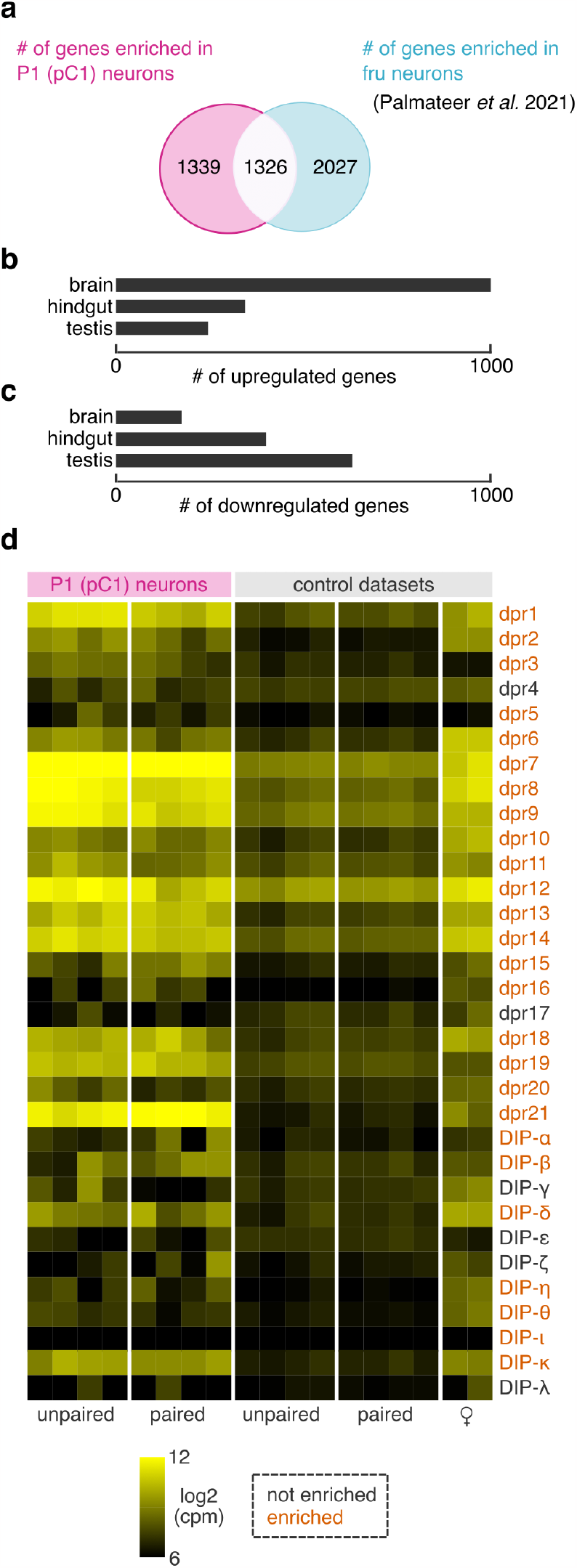
Gene expression differences between P1 (pC1) and fruitless+ neurons. **(A)** Venn diagram of 1,326 genes that are upregulated in both P1 (pC1) and fruitless+ neuron transcriptome datasets (60). **(B)** Number of the 1,326 shared genes that are upregulated in brain, hindgut, and testis-derived tissue according to FlyAtlas AffyCall. (C) Number of the 1,326 shared genes that are downregulated in brain, hindgut, and testis-derived tissue. **(D)** Expression pattern of dpr/DIP genes in P1 (pC1) and control datasets. Each row denotes a single dpr/DIP gene. Each column denotes an independent dataset (as in heatmap in Figure 1). Genes enriched in P1 (pC1) are annotated in orange.

IgSF-encoding genes, which include DPRs and their interacting DIPs, comprise cell surface molecules critical for establishing synaptic partners and differentiating cell identity (61). These genes are regulated by male-specific fruitless (FruM) (62) and at least one dpr-encoding gene, *dpr1*, contributes directly to male courtship behavior (63). In fruitless+ neurons, *dprs* are widely expressed and *DIPs* are more restricted, with many cells expressing a unique blend of each (62). These genes are also sexually dimorphic; for example, *dpr7, dpr18, dpr21, DIP-gamma, DIP-theta*, and *DIP-zeta* show a male-biased pattern of expression in fruitless+ neurons (48). Given the role of IgSF-encoding genes in cell-cell communication and cell identity, we profiled the expression of all 21 *dprs* and 11 *DIPs* in P1 (pC1) 4D). We find that most *dprs* are differentially-expressed in P1 (pC1), with the exception of *dpr4* and *dpr17*. Most *DIPs* were also enriched in P1 (pC1), except for *DIP-gamma, -epsilon, zeta*, and *lambda*. Identifying the unique combination of cell surface molecules, including dprs and DIPs, will contribute to our understanding of P1 (pC1)’s morphology and physiology.

### Courtship-related changes in P1 (pC1) gene expression

Transcriptomes are not static catalogs— they dynamically change with cell functionality and physiology. We therefore hypothesized that courtship interactions, which dynamically alter P1 (pC1) neuronal firing, will significantly impact P1 (pC1)’s transcriptome. To test this, we paired singly-housed male flies with conspecific female flies and allowed them to court for 30 minutes before we harvested P1 (pC1) neurons. Transcriptomic data from this “paired” group of male flies was compared to an “unpaired” group, which were similarly handled but were not provided with a courtship partner. Comparing these two datasets revealed 322 differentially-expressed genes, with 88 genes currently unidentified (CG names). GO enrichment analysis of the 322 genes revealed terms for “cytoplasmic translation” (21 genes encoding ribosomal proteins, p-value = 5.11e9) and “carbohydrate metabolic process” (20 genes, p-value = 2.53e-2), suggesting that many genes contribute to translation-related processes that are linked to the changing activity patterns of P1 (pC1).

We expect that the molecular mileu of P1 (pC1) neurons from paired male flies will comprise both up- and down-regulate genes compared to unpaired flies. We find that 233 of the differentially expressed genes were upregulated, of which we highlight the 100 most upregulated below (Figure 5). Two upregulated genes (*Arr1* and *Arr2*) encode arrestin proteins that are critical for wildtype olfactory processing (64). In addition, several genes encode putative enzymes that may contribute directly to behavior; for example, *Galt* regulates locomotion (65). Other upregulated genes likely operate in the same genetic pathways; a gene network analysis (38) reveals that two upregulated genes, *Impl2* and *fit*, interact with the insulin receptor-encoding gene *InR*, which is expected to be expressed in P1 (pC1) neurons (33). All three genes contribute to feeding control in flies (33, 66, 67) and potentially act in concert to regulate the balance of competing motivational drives.

**Figure 5.**
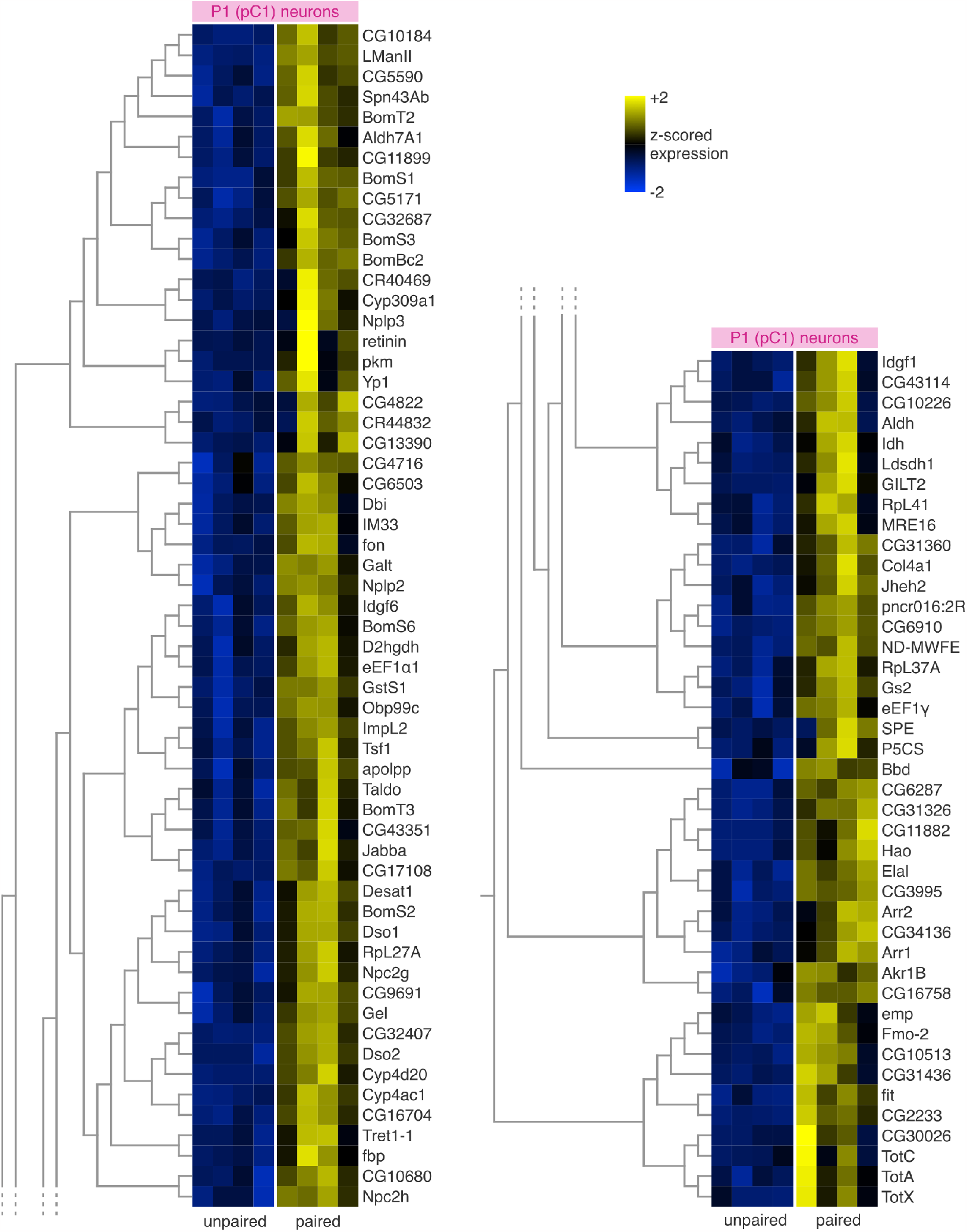
Expression of genes that are upregulated in paired male flies. The top 100 P1 (pC1) genes with higher expression in paired male flies compared to unpaired male flies. Each row denotes a single gene and each column denotes an independent dataset (as in heatmap in Figure 1). Gene expression was z-scored for each gene and genes were hierarchically clustered based on their expression pattern across animals.

Among the top 100 upregulated genes, we discovered at least 21 genes that contribute to the innate immune response in flies. These genes include *Nplp2, Col4a1, IM33, Obp99c, Bbd, SPE, GILT2, Tsf1, Jabba, Spn43Ab*, 2 Daisho genes (*Dso1* and *2*), 3 Turandot genes (*TotA, C*, and *X*), and 6 Bomanin genes (*BomBc2, S1, S2, S6, T2*, and *T3*). The Bomanins, Daishos, *Bbd*, and *SPE* are part of the Toll signaling pathway (68–70), while the Turandot genes are induced by the JAK/STAT pathway (71). We were surprised to find that more than one-fifth of the top 100 upregulated genes contributed to the immune response. We investigated this further and found support for this finding in other studies (see below).

Only ∼ 28% of differentially-expressed genes were downregulated in paired male flies (Figure 6). A closer inspection revealed that 32 of these 89 genes contribute to cell signaling (GO:0023052, p-value = 6.89e-6) and communication (GO:0007154, p-value = 1.33e-5), suggesting that courtship interactions tighten P1 (pC1) functionality by dampening expression of key signaling and synaptic genes. Some of these genes impact diverse behaviors, including locomotion: *DopEcR, Fife, Toll-6, FMRFaR* (72–75), learning and memory: *radish* (76), exercise-induced endurance: *Octbeta3R* (77), circadian rhythm: *kairos* (78), and courtship: *Ddc* (79).

**Figure 6.**
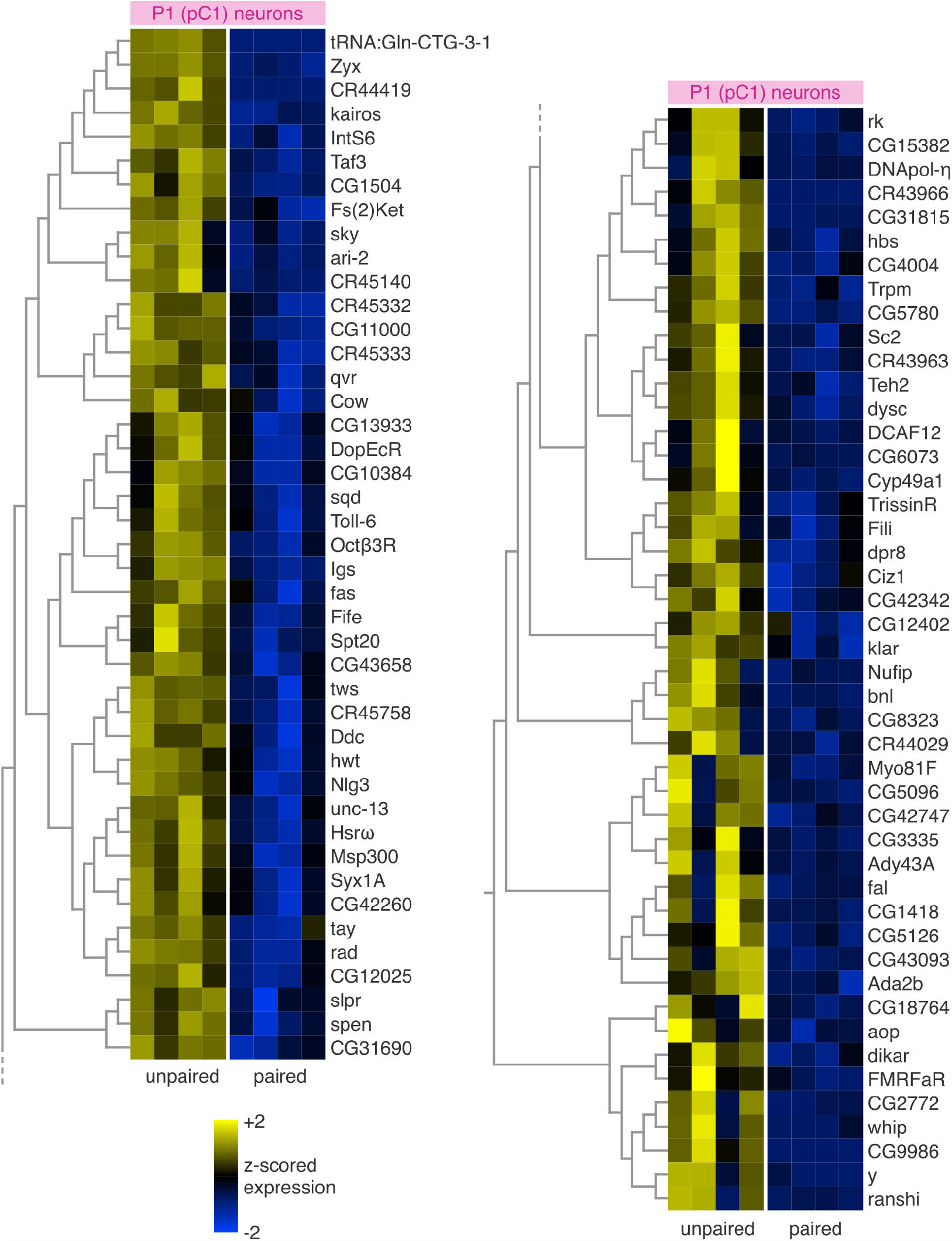
Expression of genes that are downregulated in paired male flies. The top 89 P1 (pC1) genes with lower expression in paired male flies compared to unpaired male flies. Each row denotes a single gene and each column denotes an independent dataset (as in heatmap in Figure 1). Gene expression was z-scored for each gene and genes were hierarchically clustered based on their expression pattern across animals.

While male-female contact via the foreleg tarsi increases the activity of P1 (pC1) neurons, these neurons are not persistently active (22). We find that the dynamic, courtship-dependent activity patterns of P1 (pC1) are marked by changes in many upregulated genes, many of which contribute to the immune response, and fewer downregulated genes, many of which contribute to synapse physiology and diverse behaviors. These genes likely act in concert to shift the functional characteristics of P1 (pC1) neurons as flies ramp up or down their courtship behaviors.

### Courtship-related differential expression of immuneresponse genes

We identified 322 differentially expressed genes in P1 (pC1) neurons harvested from paired male flies. Are these genes specific to P1 (pC1)? To test this we compared our list of 322 genes with two other datasets: Winbush *et al*. (35) and Jones *et al*. (80) (Figure 7). Briefly, Winbush *et al*. assayed gene expression in whole fly heads 24 hours after male flies were paired with a mated female, or left unpaired, for 10-12 hours. Through this, they identified 91 differentially-expressed genes. Jones *et al*. performed similar experiments but assayed gene expression 1 hour after pairing for 7 hours. Through this, they identified 322 differentially-expressed genes; it is by sheer coincidence that our study also identified 322 genes. Keep in mind that in our study, we paired male flies for 30 minutes, and then harvested P1 (pC1) neurons within 30 minutes. By comparing these three datasets, we revealed sets of genes that change their expression with courtship-related interactions (Figure 7). These overlapping genes were robust to technical (e.g. handling, *etc*.) and biological (e.g. genetic strain, *etc*.) differences between the studies. More importantly, because Winbush *et al*. and Jones *et al*. sequenced RNA from whole fly heads, it is unlikely that these overlapping genes comprise a P1 (pC1)-specific gene response. By extension, non-overlapping genes (the majority of genes) may act more specifically in P1 (pC1) (Figures 5 and 6). Out of the 322 genes in our study, we identified 54 genes that overlapped with Winbush *et al*. (n = 26), Jones *et al*. (n = 19), or both datasets (n = 9) (Figure 7). Of these genes, 12 are unidentified (CG names) and 5 encode ribosomal proteins. The vast majority of overlapping genes (51 of 54 genes) were upregulated in P1 (pC1) neurons, more than would be expected by chance (Fisher’s exact test, p < .01). Of these, 29 genes appeared in our top 100 most upregulated genes list (harken back to Figure 5). In contrast, only 3 overlapping genes (*CG12025, Toll-6*, and *Msp300*) appeared in our downregulated list. This difference could be due to many factors, one of which is that the upregulated overlapping genes constitute a generic neuronal response to courtship pairing. The downregulated genes, on the other hand, could reflect P1 (pC1)-specific regulatory responses that are not captured at the level of the whole head. Indeed, as we have shown above, many genes downregulated in P1 (pC1) are involved in cell-specific processes such as synapse physiology, cellular communication, and behavioral control.

**Figure 7.**
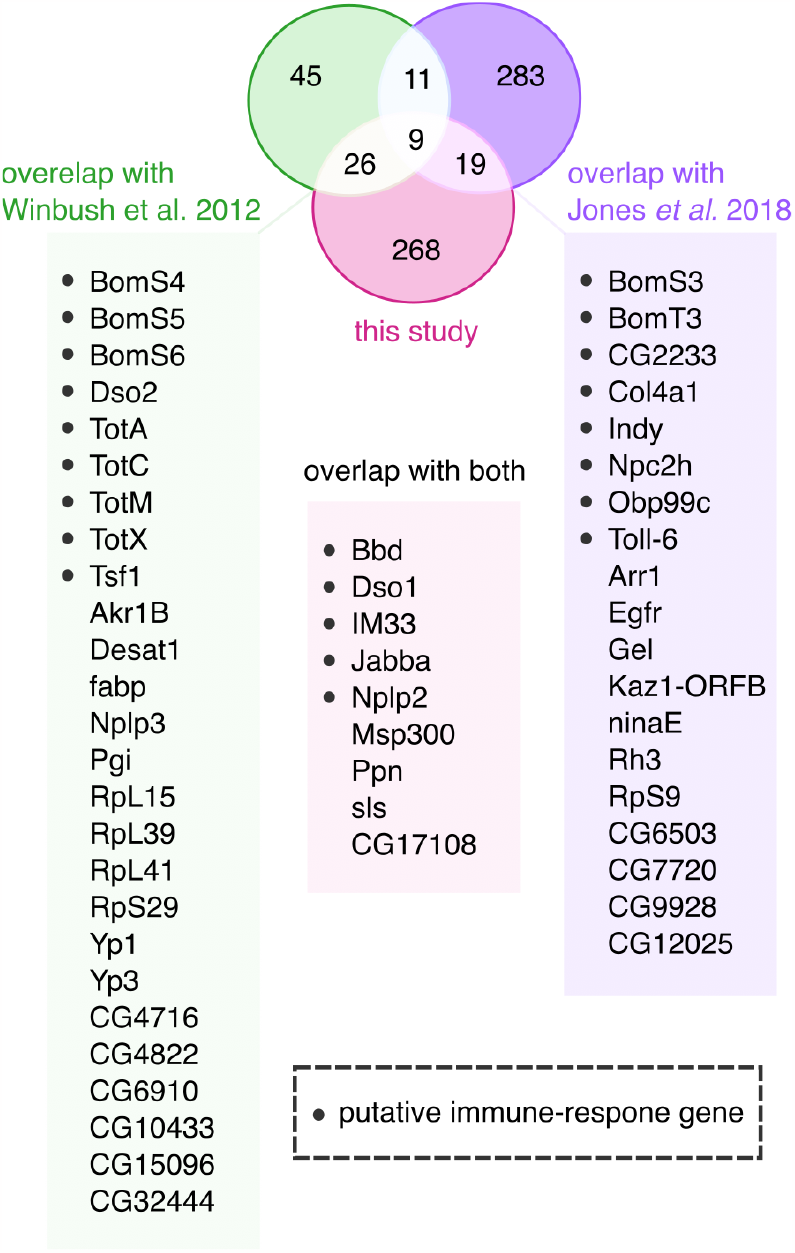
Differential gene expression associated with courtship. Venn diagram of genes identified in this study with differential expression associated with courtship pairing, compared with two studies that identified genes with differential expression in the head following courtship pairing (35, 80). Genes with a putative role in the immune-response are highlighted.

Approximately 41% (22 out of 54) of the overlapping genes contribute to the immune response. Out of these genes, 9 overlapped with Winbush *et al*., 8 with Jones *et al*., and 5 with both studies (Figure 7). These genes include, among others, 4 Turandots (*TotA, C, M*, and *X*), 2 Daishos (*Dso1* and *Dso2*), 5 Bomanins (*BomS3, S4, S5, S6*, and *T3*), *Toll-6*, and Bombardier (*Bbd*), which interacts with Bomanins in the Toll-mediated immune response (81). Apart from these gene families, we also find interactions between three other overlapping immuneresponse genes (*Obp99c, Tsf1*, and *Nplp2*); these three genes also interact with *Apolpp*, which is in the top 100 most upregulated gene list (Figure 5). Overall, these findings link male fly courtship behavior and the innate immune response at the neural level.

## Discussion

Single-cell RNA sequencing has uncovered the heterogeneous and complex transcriptional profiles of large neuronal populations, revealing distinct subpopulations (49, 82, 83), sex dimorphisms (48), and state-dependent gene-expression patterns (84– 88) in the fly brain. Our work complements these approaches by profiling a much smaller population (10-12 neurons per fly) and therefore boosts resolution while capturing fine-scale heterogeneity (36) and dynamic changes. Moreover, the population we target, P1 (pC1), is critical for many fly behaviors, including driving a long-lasting motivational state, and the diverse functions of P1 (pC1) neurons are tied to the genes they express. Here we uncover these genes via RNA-sequencing, revealing sets of genes that encode receptors, synaptic signaling molecules, and cell surface proteins. Approximately half of P1 (pC1)-expressed genes are not shared with the majority of neurons in their superset population, the ∼ 2,000 fruitless+ neurons. About 300 genes dynamically change expression when male flies are paired with female conspecifics, compared to unpaired male flies. Among these genes, many immune-response genes are upregulated while many synaptic signaling genes, many of which regulate diverse behaviors, are downregulated. All together, these results unravel the dynamic, molecular landscape of an important set of higher-order neurons. This work fills an important gap and dovetails nicely with recent single-cell RNA sequencing studies of larger populations of neurons.

Although our study provides valuable insights, it has some limitations. Firstly, we only profiled a subset of the approximately 40 P1(pC1) neurons, which are heterogeneous and composed of at least two genetically-defined subtypes (9). Sparsifying P1(pC1) neurons through genetic intersection of multiple markers can help resolve this issue, and our dataset can serve as a resource for potential gene selection. Secondly, it is difficult to distinguish gene expression changes due to courtship behaviors per se from those resulting from the sensory outcomes of social interactions. This limitation is inherent to all studies that use social pairing as a test variable. Additionally, we only harvested neurons and sequenced RNA at one timepoint, shortly after social pairing, which places a conservative cap on the number of overlapping genes we found when we compared our data with two previous studies that waited many hours before collecting tissue (Figure 7). In these comparisons, we may have also missed overlapping genes due to technical differences in our analyses pipelines, including differences in cutoff thresholds for example (35, 80). Despite this, we identified more than 300 genes with differential expression after social pairing and more than 50 overlapping genes, indicating that we sequenced RNA at a transcriptionally meaningful timepoint. Lastly, because we were interested in courtship-related changes in gene expressed, we tested male flies with mated female flies in order to prevent copulation. However, P1(pC1)’s transcriptomic profile may differ when male flies court unmated female flies or fight other male flies. Future experiments with P1 (pC1) RNA sequencing from these conditions can be compared directly to our dataset.

One surprising finding of our study was uncovering that immune-response genes are upregulated in paired male flies compared to unpaired. Many of these gene responses are likely not specific to P1 (pC1) as we find similar results in two other studies (35, 80). This raises the question: why are immuneresponse genes upregulated following courtship interactions? One possibility is that courtship is inherently risky, requiring close contact with individuals harboring potential pathogens (bacteria, fungi, viruses, and parasites) (89) and these commensal microorganisms can directly impact neuronal function and behavior (90–94). The risk, and impact, of infection may have driven flies to evolve countermeasures. For example, female flies ramp up immune-response genes (including Turandot genes we find upregulated in our study) after hearing courtship song signals, even in the absence of a male fly (95). This suggests that the mere potential of courtship interactions suffices to kick the immune system into high gear, providing an adaptive strategy for protecting against the transmission of pathogens during close-contact social encounters (96). This phenomenon is not specific to courtship interactions; co-housed male flies upregulate immune-response genes compared to socially isolated flies (97). This socially-driven immune response likely dampens the impact of socially-transmitted infections which themselves may alter behavior (98, 99). Our results therefore highlight an important two-way street between social behaviors and the immune system and future studies may reveal the impact of immune-response genes on P1 (pC1) functionality and the behaviors these neurons control.

## Methods

### Flies

For all experiments, we used singly-housed 3-7 day old male flies of the genotype NP2631-GAL4, fruFLP, UAS-mCD8-GFP. This line labels 14 neurons per hemibrain in the P1 cluster (11). Male flies were either placed with mated female flies for 30 minutes (“paired” group, male flies courted avidly but did not copulate during this period), or placed alone for 30 minutes (“unpaired” group), in a small circular arena. We then harvested P1 (pC1) neurons for RNA sequencing.

### Harvesting P1 (pC1) neurons for RNA sequencing

P1 (pC1) neurons, as defined by the intersection of NP2631GAL4 and fruFLP, were harvested as described by Crocker *et al*. 2016 (36). Briefly, we harvested 10-12 GFP+ cells per fly into 0.5ul nuclease free water in the pipette tip and then the tip was broken into a 96 well PCR tube containing RNAse inhibitors and buffer as described by Clontech’s ultra low HV SMARTer Ultra Low RNAseq kit (Catalog #634823). We then used the Clontech Ultra-Low high volume SMARTer RNAseq kit for amplification. We performed 15 rounds of PCR amplification using the Clontech SMARTer Ultra low RNAseq Kit. The Princeton University Sequencing Core prepared cDNA and libraries. Samples were fragmented into 200bp fragments then libraries were made using IntegenX’s Apollo 324 automated library prep system. We used the PrepX Illumina DNA library prep kit/PrepX CHIPseq kit (WaferGen Biosystems Inc) and ran 17-22 PCR cycles. We then barcoded samples (Bio Scientific) and ran them on the Illumina HiSeq2500 with a depth of 30 million reads.

### Analysis of Sequencing Reads

We used the Galaxy web platform to map and analyze our reads and the public server at usegalaxy.org to analyze the data (100). We used FastQC (101) to evaluate reads and to remove samples of poor quality; we excluded one male fly because read count was an order of magnitude smaller than other flies. All mapping was performed using the Galaxy server (v. 21.01) running Hisat2 (Galaxy Version 2.1.0+galaxy7) (102), FeatureCounts (Galaxy Version 2.0.1) (103), and Deseq2 (Galaxy Version 2.11.40.6+galaxy1) (104). We used the UCSC Reference sequence Dm6 and the GTF files (Ensembl – BDGP6.87). We ran Hisat2 with the following parameters: single-end, unstranded, and default settings (except for a GTF file that was used for transcript assembly). The aligned SAM/BAM files were processed using Featurecounts with default settings, except we used Ensembl BDGP6.87 GTF file and output for DESeq2 and gene length file. FeatureCounts output files and raw read files are publicly available (GEO with accession GSE223552). Gene counts were normalized using DESeq2 followed by a regularized log transformation. We used DESeq2 to determine differential expression, using the following settings: for factors we used “cell-type” (P1 (pC1), female brain, or whole head) or “condition” (unpaired vs. paired) and then ran pairwise comparisons to generate normalized tables. For size estimation we used the standard median ratio, with a parametric fit type, and we filtered outliers with Cook’s distance cutoff. We then used Qlucore Omics Explorer to generate heatmaps and volcano plots.

For our control datasets comprised transcriptomes of 2 female whole brains and 8 male whole head samples. Female brain samples were taken from Crocker et al. 2016 (36) (GEO Accession GSE74989) (36) and whole head samples were pulled from Winbush *et al*. 2012 (35) following remapping of SRA data sets (GEO Accession Info GSE 117217). We used the following datasets in our analyses: GSM3278900_SRX170947, GSM3278901_SRX170948, GSM3278902_SRX170949, GSM3278903_SRX170950, GSM3278904_SRX170951, GSM3278905_SRX170952, GSM3278906_SRX170953, GSM3278907_SRX170954.

### Gene Ontology and Network Analysis

We used FlyMine.org (38) to run gene ontology analyses and to test for gene network interactions.

### Gene expression comparison to other studies

For Figure 7, we pulled data whole head transcriptome data from Winbush et al. 2012 and Jones et al. 2018. In Winbush *et al*., we treated “naive” male flies as “unpaired”, and “trained” flies as “paired”. These flies were tested for 12-14 hours in similarly sized chambers to ours (1cm space) and then whole heads were harvested and RNA sequenced 24 hours after. In Jones et al. 2018 (80), we used data from the 1hr timepoint because this aligned closely with our protocol for harvesting P1 (pC1) neurons. To compare fruitless+ neuron gene expression with P1 (pC1) gene expression, we pulled the 10-12 day old male dataset from Palmateer *et al*. 2021 (60).

## Data Availability

Raw data is publicly available (GEO accession GSE223552). Processed gene count Txt files are also available under the same GEO accession GSE22352.

## AUTHOR CONTRIBUTIONS

OMA— Conceptualization, Validation, Visualization, Writing – original draft, Writing – review and editing. AC— Conceptualization, Data curation, Formal Analysis, Investigation, Methodology, Validation, Visualization, Writing – review and editing. MM— Conceptualization, Funding acquisition, Methodology, Project administration, Resources, Supervision, Writing – review and editing.

## ACKNOWLEDGEMENTS

We thank Drs. Mike Crickmore, Maxi Pitsch, Ethan Glantz, and Z. Yan Wang for critical feedback on the manuscript. OMA was supported by the Burroughs Wellcome Fund Postdoctoral Enrichment Program, the BRAINS Fellowship, the Simons Collaboration on the Global Brain Bridge to Independence Award, and the Princeton Presidential Postdoctoral Fellowship. AC was supported by an Institution Development Award (IDeA) from the National Institute of General Medicine (NIGMS) of the National Institutes of Health grant number P20GM103449, NIH/NIGMS F32 grant number GM105370-01, and an R15 from NIGMS grant number R15GM132937. This work was supported by an NIH New Innovator Award, NSF CAREER Award and McKnight Foundation Scholar Award to MM. Drawings and schematics in Figure 1 are provided by Julie Johnson (Life Science Studios) and via biorender.com. We acknowledge that this research was conducted on the unceded territories of the Nanticoke Lenni-Lenape Tribal Nation.

## Supplement

**Figure S1.**
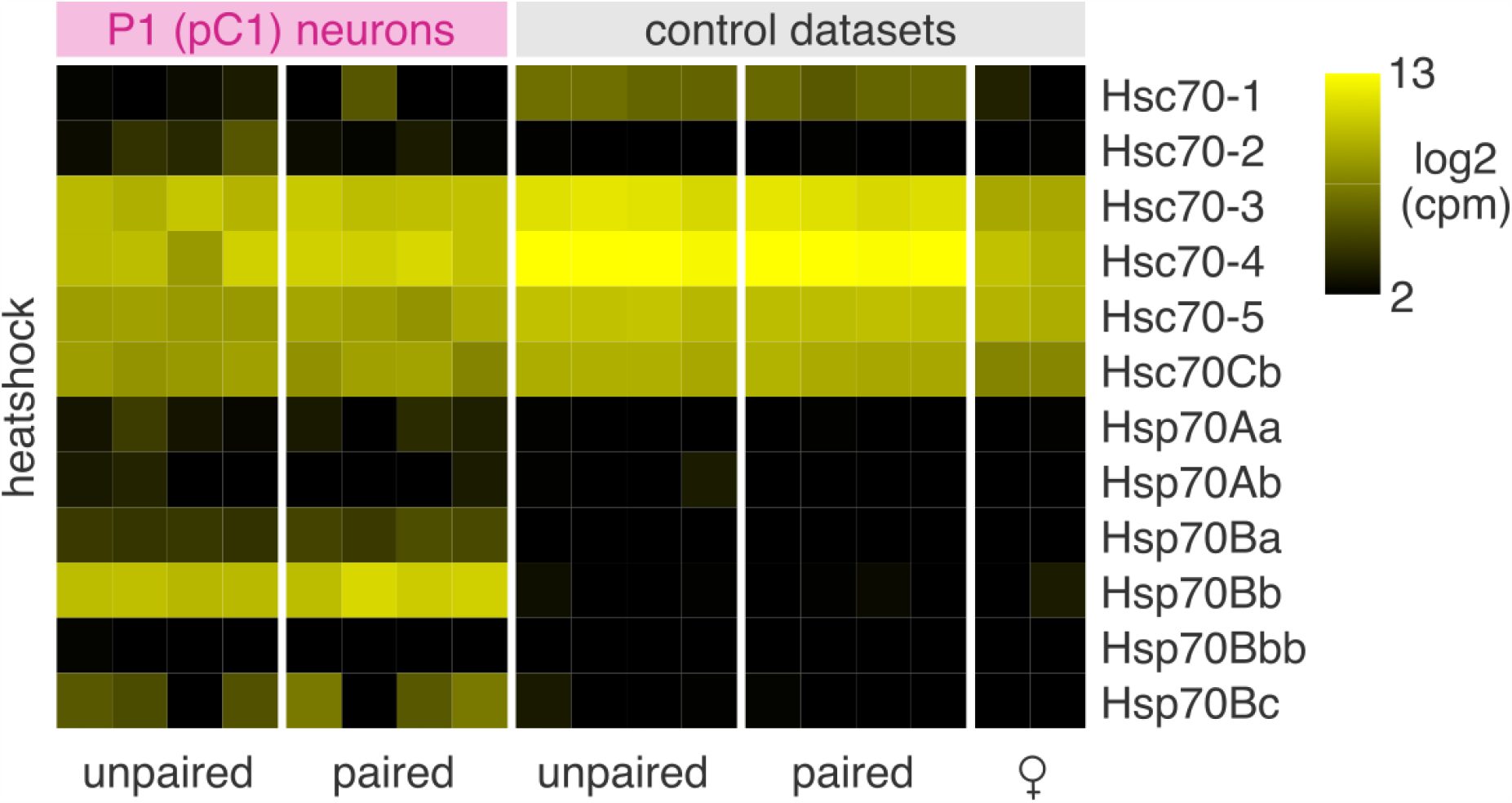
Heatshock protein-encoding gene expression of P1 (pC1) neurons. Each row denotes a single gene and each column denotes an independent dataset (as in heatmap in Figure 1).

**Figure S2.**
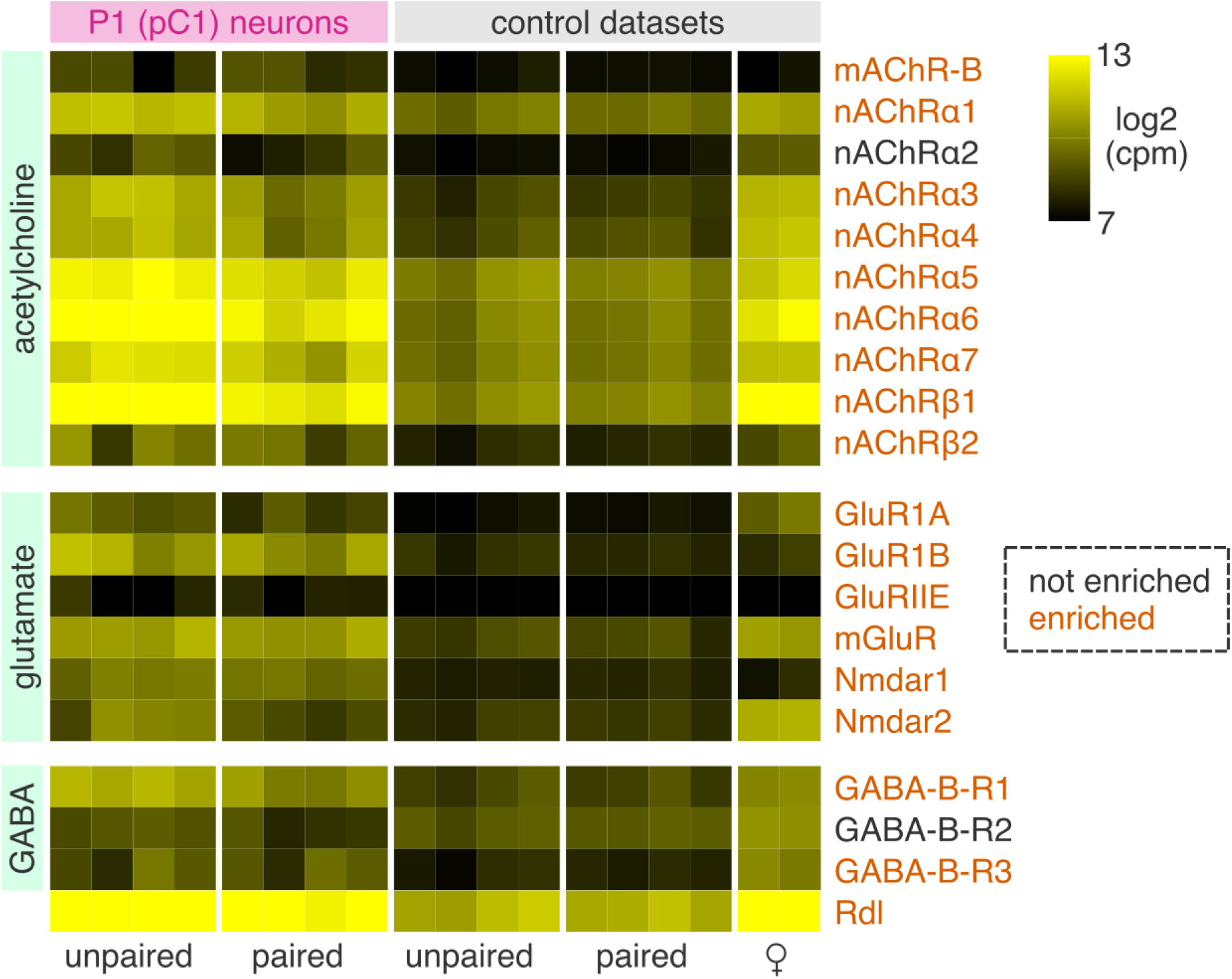
Canonical neurotransmitter receptor genes. Each row denotes a single receptor gene and each column denotes an independent dataset (as in heatmap in Figure 1).

**Figure S3.**
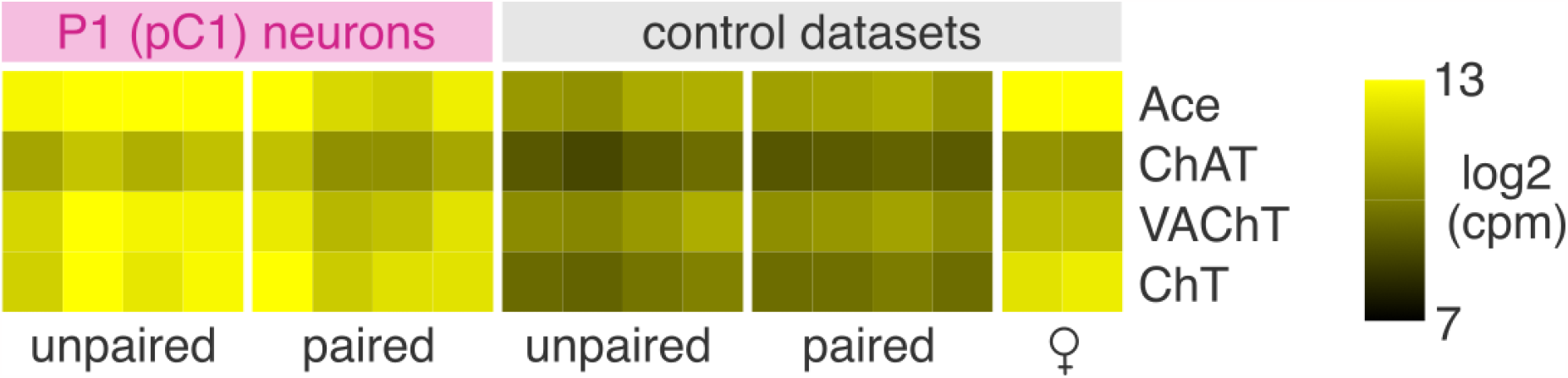
P1 (pC1) neurons are cholinergic. Each row denotes a single gene and each column denotes an independent dataset (as in heatmap in Figure 1).

